# Lack of pathogenic involvement of CCL4 and its receptor CCR5 in arthritogenic alphavirus disease

**DOI:** 10.1101/2024.07.31.606106

**Authors:** Muddassar Hameed, Norman A. Solomon, James Weger-Lucarelli

## Abstract

Arthritogenic alphaviruses, including chikungunya virus (CHIKV), Mayaro virus (MAYV), Ross River virus (RRV), and O’nyong nyong virus (ONNV) are emerging and reemerging viruses that cause disease characterized by fever, rash, and incapacitating joint swelling. Alphavirus infection induces robust immune responses in infected hosts, leading to the upregulation of several cytokines and chemokines, including chemokine C ligand 4 (CCL4). CCL4 is a chemoattractant for immune cells such as T cells, natural killer cells, monocytes/macrophages, and dendritic cells, recruiting these cells to the site of infection, stimulating the release of proinflammatory mediators, and inducing T cell differentiation. CCL4 has been found at high levels in both the acute and chronic phases of chikungunya disease; however, the role of CCL4 in arthritogenic alphavirus disease development remains unexplored. Here, we tested the effect of CCL4 on MAYV infection in mice through antibody depletion and treatment with recombinant mouse CCL4. We observed no differences in mice depleted of CCL4 or treated with recombinant CCL4 in terms of disease progression such as weight loss and footpad swelling or the development of viremia. CCL4 uses the G protein-coupled receptor C-C chemokine receptor type 5 (CCR5). To determine whether CCR5 deficiency would alter disease outcomes or virus replication in mice, we inoculated CCR5 knockout (CCR5^-/-^) mice with MAYV and observed no effect on disease development and immune cell profile of blood and footpads between CCR5^-/-^ and wild type mice. These studies failed to identify a clear role for CCL4 or its receptor CCR5 in MAYV infection.

## Introduction

Arthritogenic alphaviruses, including chikungunya virus (CHIKV), Ross River virus (RRV), O’nyong nyong virus (ONNV), and Mayaro virus (MAYV) are significant public health threats (1, 2). Arthritogenic alphaviruses are found worldwide: CHIKV is endemic to Africa, Southeast Asia, and, more recently, the Caribbean and South America, while RRV, ONNV, and MAYV circulate in Australia, Africa, and South America, respectively (3). These viruses cause acute and chronic disease characterized by high fever, rash, and muscle and joint inflammation, which can persist for years in roughly half of affected patients (4–9). There are no specific drugs available for the treatment of alphavirus arthritis except the use of anti-inflammatory drugs for symptomatic relief (10, 11). Thus, there is an urgent need to understand the immune mechanisms that control arthritogenic alphavirus disease outcomes in order to develop therapeutics.

During arthritogenic alphavirus infection, the virus replicates in fibroblasts, muscle satellite cells, macrophages, and other cells, initiating inflammatory responses (12, 13). This results in the influx of monocytes, macrophages, natural killer cells, neutrophils, and T and B lymphocytes to the infection site, leading to tissue damage and expression of proinflammatory cytokines and chemokines, including chemokine C ligand 4 (CCL4), further exacerbating inflammation (14–16). Previous studies have shown that CCL4 (also known as macrophage inflammatory protein-1β or MIP-1β) is upregulated in acute and chronic arthritogenic alphavirus-infected humans (17–19) and mice (20–22). CCL4 is involved in orchestrating immune cell movement and activation during infection or inflammation (23–26). CCL4 binds to CC motif chemokine receptor 5 (CCR5) (27), acting as a chemoattractant for T cells, monocytes, natural killer cells, and dendritic cells, which play an important role in arthritogenic alphavirus pathogenesis (7, 13, 14, 24, 26, 28). Therefore, understanding CCL4’s effect on disease outcomes during arthritogenic alphavirus infection is critical to elucidate its contribution to pathogenesis and, thus, its potential to be targeted for the development of therapeutics.

In this study, we investigated the contribution of CCL4 in the development of MAYV-induced arthritis and disease. We blocked CCL4 in wild-type (WT) mice using an anti-CCL4 monoclonal antibody (mAb) followed by MAYV infection and observed no effect on disease severity. We also inoculated mice with recombinant mouse CCL4 and then infected with MAYV, which similarly showed no effect on disease outcome. Finally, we infected mice deficient in CCR5 with MAYV and observed no differences in disease severity or immune cell profiles at different timepoints of infection compared to WT mice. These studies failed to identify a clear role for CCL4 or its receptor CCR5 in MAYV infection.

## Materials and Methods

### Ethics statement

All experiments were conducted with the approval of Virginia Tech’s Institutional Animal Care & Use Committee (IACUC) under protocol number 21-041. Experiments using MAYV were performed in a BSL-2 facility in compliance with CDC and NIH guidelines and with approval from the Institutional Biosafety Committee (IBC) at Virginia Tech.

### Mice

C57BL/6J mice (strain #000664) and chemokine C-C motif receptor type 5 knock-out (CCR5^-/-^) mice (B6.129P2-CCR5^tmKuz/^J; strain #005427) were purchased from The Jackson Laboratory at 6-8 weeks of age. CCR5^-/-^ mice were bred at Virginia Tech, and 6–8-week-old mice were used for experiments. Mice were kept in groups of five animals per cage at ambient room temperature with *ad libitum* supply of food and water.

### Cell culture and viruses

Vero cells were obtained from the American Type Culture Collection (ATCC; Manassas, VA) and grown in Dulbecco’s Modified Eagle’s Medium (DMEM, Gibco) with 5% fetal bovine serum (FBS, Genesee), 1 mg/mL gentamicin (Thermo Fisher), 1% non-essential amino acids (NEAA, Sigma) and 25 mM HEPES buffer (Genesee) at 37°C with 5% CO_2_. MAYV strain TRVL 4675 was derived from an infectious clone (29, 30). Virus titers were determined by plaque assay as previously described (31).

### Mouse infections

Mice were injected through both hind footpads with 10^4^ PFU of MAYV in 50 μL viral diluent in each foot (32). All virus dilutions were made in RPMI-1640 media with 10 mM HEPES and 1% FBS. Mice were monitored for disease development following infection through daily weighing, and footpad swelling was measured using a digital caliper. Blood was collected via submandibular bleed for serum isolation to determine viremia and cytokine levels. At six-, seven-, or twenty-one-days post-infection (dpi), mice were euthanized, and blood and footpads were collected to isolate serum and immune cells. Leukocytes were isolated from blood and footpads to perform flow cytometry.

### Luminex assay and ELISA

We quantified CCL3, CCL4, and CCL5 levels in the serum of mock- and MAYV-infected animals using the mouse Luminex XL cytokine assay (bio-techne) and the CCL4/MIP-1 beta DuoSet ELISA kit (Catalog# DY451-05, R&D Systems) according to the manufacturer’s instructions. The standard curve was generated using the optical density values of the standards, which were used to calculate the cytokine levels in each sample.

### CCL4 antibody and cytokine treatment

For *in vivo* CCL4 neutralization, 6–8-week-old C57BL/6J mice were intravenously inoculated through the retro-orbital sinus with 20 μg/mouse of Rat IgG2a kappa isotype control (Cat. No. 50-112-9680, Fisher Scientific) or an anti-mouse CCL4 mAb (Cat. No. PIMA523742, clone 46907, Fisher Scientific) at -1, 1, 3, and 5 dpi, as previously described (33). Antibodies were diluted in phosphate-buffered saline (PBS, Genesee) and administered in a volume of 100 μL. For gain of function studies, 6–8-week-old C57BL/6J mice were administered PBS or recombinant mouse CCL4 (400 ng/mouse, Cat. No. 554602, BioLegend) intraperitoneally at -3, - 1, 1, 3, and 5 dpi (34). Our rationale for the dosage and schedule of CCL4 treatment was based on a previous study that showed that a single dose of 500 ng CCL4 increased macrophages two-fold in BALF in mice with *S. aureus* infection (34). Other studies have used recombinant cytokines for multiple days and observed a significant impact on immune cell mobilization (35–37). Thus, we reasoned those multiple injections of CCL4 prior to and after infection would elicit a more robust effect on immune cell infiltration to the infected tissues. Mice were then infected with 10^4^ PFU of MAYV in both hind feet and monitored for disease development until 7 dpi.

### Mouse blood and footpad immune cell isolation

Mouse blood leukocytes were isolated using Mono-Poly resolving medium (M-P M; MP Bio, Cat. No. 091698049) according to the manufacturer’s instructions. Briefly, blood was mixed with an equal volume of PBS and layered slowly onto M-P M followed by centrifugation at 300 × *g* for 30 min in a swinging bucket rotor at room temperature (20–25°C). We collected cell layers between the plasma and M-P M to isolate leukocytes and added them to a 15 mL conical tube containing 10 mL cold 10% FBS containing RPMI-1640 (RPMI-10). Cells were spun at 500 × *g* for 5 min at 4°C and used for flow cytometry. We isolated footpad immune cells as we previously described (38). Briefly, footpads were collected above the ankle, deskinned, and transferred to digestion media [RPMI-10, 2.5 mg/mL Collagenase I (Cat. No. LS004196, Worthington Biochemical Corporation), 17 μg/mL DNase I (Cat. No. LS006333, Worthington Biochemical Corporation)], incubated for 2 hours at 37°C, and filtered through a 70 μM cell strainer followed by washing with RPMI-10.

### Flow cytometry

Single cell suspensions were washed with PBS and resuspended in 100 μL Zombie aqua cell viability dye solution (1:400 prepared in PBS, Cat. No. 423101, BioLegend) and incubated at room temperature for 15-30 minutes. 200 μL flow cytometry staining (FACS) buffer (PBS containing 2% FBS) was added and centrifuged at 500 × *g* for 5 min at 4°C. The resulting cell pellet was resuspended in FACS buffer with 0.5 mg/mL rat anti-mouse CD16/CD32 Fc block (Cat. No. 553142, BD Biosciences) and incubated for 15 min on ice to block Fc receptors. For extracellular staining, a combined antibody solution was prepared in FACS buffer with fluorophore-conjugated antibodies: anti-mouse Alexa fluor 700 CD45 (Cat. No. 103128, BioLegend), anti-mouse PerCP/Cyanine 5.5 CD11b (Cat. No. 101227, BioLegend), anti-mouse brilliant violet 421 F4/80 (Cat. No. 123131, BioLegend), anti-mouse APC Ly6G (Cat. No. 127614, BioLegend), anti-mouse PE Ly6C (Cat. No. 128007, BioLegend), anti-mouse PE-Dazzle 594 MHC II (Cat. No. 107647, BioLegend), anti-mouse PE CD3 (Cat. No. 100206, BioLegend), anti-mouse PerCP/Cyanine 5.5 CD4 (Cat. No. 116012, BioLegend), anti-mouse FITC CD8a (Cat. No. 100706, BioLegend), anti-mouse brilliant violet 421 NK1.1 (Cat. No. 108741, BioLegend), and anti-mouse Alexa fluor 488 CD11c (Cat. No. 117311, BioLegend). 100 μL antibody cocktail was added to the single cell suspension, mixed, and incubated for 30 min on ice. Cells were washed with FACS buffer twice, and 100 μL 4% formalin (Thermo Fisher Scientific, Ref. No. 28908) was added to fix the cells. After 15 min incubation at room temperature, cells were washed with FACS buffer, resuspended in 100-200 μL PBS, and covered with aluminum foil before flow cytometry analysis. For each antibody, single color controls were run with Ultracomp ebeads (Cat. No. 01-2222-42, Thermo Fisher Scientific). The stained cells were analyzed using the FACSAria Fusion Flow cytometer (BD Biosciences).

### Statistical analysis

All statistics were performed using GraphPad Prism version 9 and data are presented as mean ± standard deviation. The statistical tests used to analyze data are described in the figure legends.

## Results

### CCL4 is upregulated in response to MAYV infection in C57BL/6J mice

CHIKV and MAYV produce disease in C57BL/6J mice similar to outcomes in humans (32). MAYV is a BSL2 virus that induces similar disease in humans and WT mice to CHIKV, allowing for safer and easier handling than CHIKV, which requires BSL3 conditions; therefore, to explore CCL4’s effect on arthritogenic alphavirus infection, we used MAYV as a model arthritogenic alphavirus (29). CCL4 is involved in inflammatory responses and is elevated in humans infected with arthritogenic alphaviruses (17–19). To assess CCL4 expression induced by MAYV, we infected mice with MAYV and collected blood at 2 and 7 days post-infection (dpi), the peak of viremia and footpad swelling, respectively (29). At 2 dpi, we found that CCL4 was significantly upregulated following MAYV infection compared to mock-infected controls (Fig. 1A). To assess whether CCL4 remained elevated later in infection, we measured levels in the blood at 7 dpi. Similarly, CCL4 levels were higher in MAYV-infected mice compared to the mock-infected group (Fig. 1B). This higher expression of CCL4 in MAYV-infected mice is consistent with reports in humans infected with MAYV or CHIKV (8, 19, 39).

**Figure 1.**
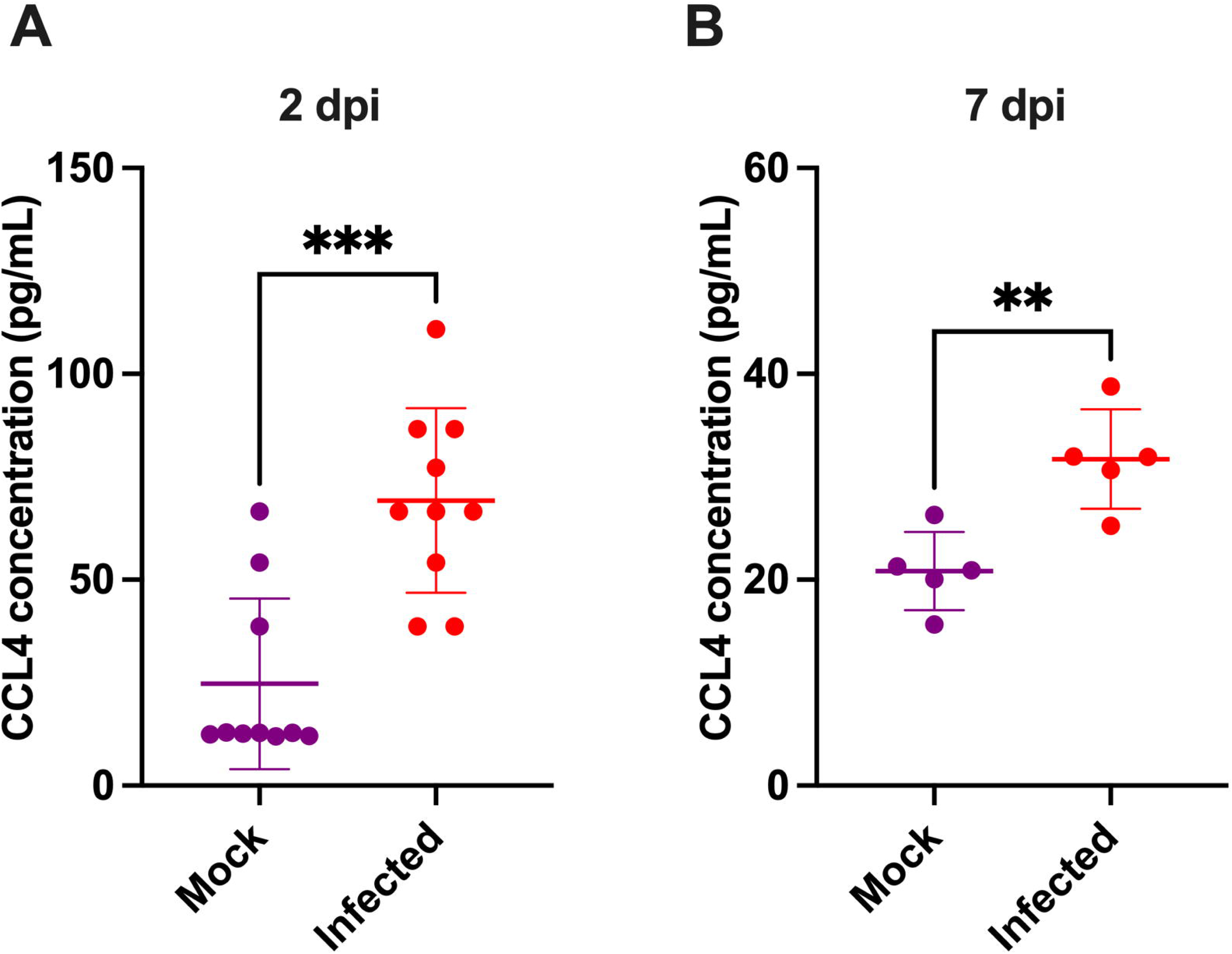
CCL4 is upregulated in response to MAYV infection in C57BL/6J mice. C57BL/6J mice were inoculated with RPMI-1640 media (mock) or 10^4^ PFU of MAYV strain TRVL 4675 in both hind feet and monitored for disease development until 7 dpi (n=5 or 10). CCL4 was measured by Luminex assay at 2 dpi (A) and ELISA at 7 dpi (B) and is presented in picograms/mL of serum. Statistical analysis was done using an unpaired t-test with Welch’s correction. The error bars represent the standard deviation, bars indicate mean values, and asterisks indicate statistical differences; **, p < 0.01; ***, p < 0.001.

### Antibody-mediated CCL4 depletion and administration of recombinant mouse CCL4 cytokine failed to alter Mayaro disease severity

CCL4 acts as a chemoattractant for different immune cells, such as monocytes, macrophages, natural killer cells, dendritic cells, and T cells, which play important roles in arthritogenic alphavirus pathogenesis (40, 41). After observing higher CCL4 expression following MAYV infection, we asked whether *in vivo* blockade of CCL4 would alter disease severity. To that end, mice were intravenously inoculated with IgG2a isotype control or anti-mouse CCL4 monoclonal antibody (mAb) followed by MAYV infection and monitored for disease development as previously described (32). Mice treated with anti-mouse CCL4 mAb showed modestly higher weight gain and less footpad swelling but showed no statistical differences up to 7 dpi compared to control mice (Fig. 2A-B). Furthermore, a similar titer of infectious virus was observed in isotype and anti-mouse CCL4 mAb treated group at 2 dpi (Fig. 2C).

**Figure 2.**
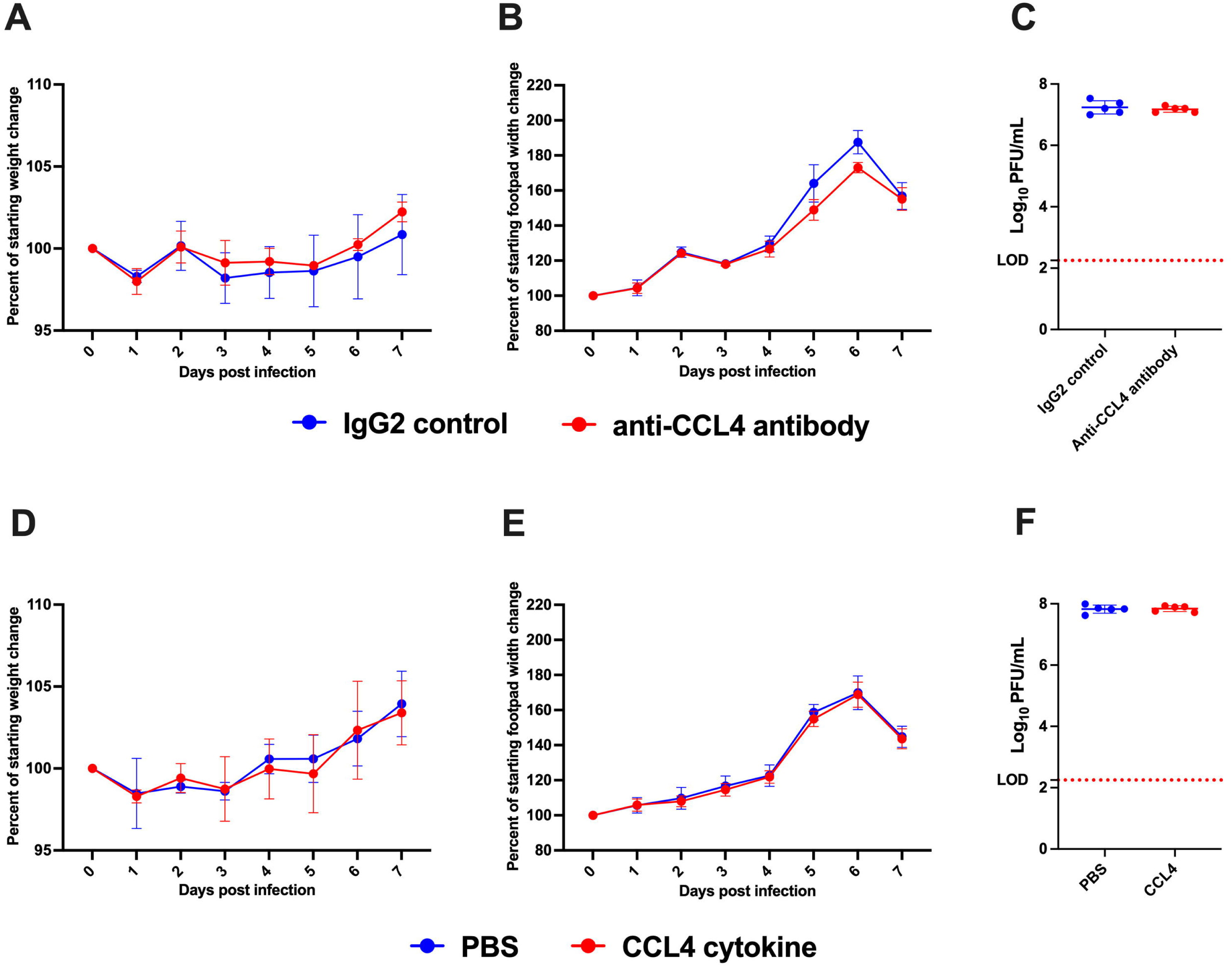
Antibody-mediated CCL4 depletion and administration of recombinant mouse CCL4 cytokine failed to alter Mayaro disease severity. C57BL/6J mice were intravenously inoculated with IgG2a isotype or anti-mouse CCL4 mAb (20 μg/mouse) at -1, 1, 3, and 5 dpi (n=5). For CCL4 cytokine treatment, C57BL/6J mice were treated with PBS or recombinant mouse CCL4 (400 ng/mouse) through the intraperitoneal route at -3, -1, 1, 3, and 5 dpi (n=5). Mice were inoculated with 10^4^ PFU of MAYV in both hind feet after the first dose of CCL4 antibody or the second dose of CCL4 cytokine and monitored for disease development until 7 dpi. A-C. Weight loss (A), footpad swelling (B), and the development of viremia (C) were determined after CCL4 depletion in C57BL/6J mice. D-E. Weight loss (D), footpad swelling (E), and virus replication (F) were measured after CCL4 cytokine treatment. Weight loss and footpad swelling were analyzed using multiple unpaired t-tests with the Holm-Sidak method for multiple comparisons, and viremia data was analyzed by unpaired t-test with Welch’s correction. The error bars represent the standard deviation, the solid line indicates mean values, and the dotted line represents the limit of detection.

Next, we tested whether CCL4 protein inoculation would impact Mayaro disease. We treated mice with PBS or recombinant mouse CCL4 at multiple timepoints before and after MAYV infection and monitored for disease development and virus replication. We found no differences in weight loss or footpad swelling up to 7 dpi (Fig. 2D-E). Likewise, no difference was seen in viral replication between PBS injected and CCL4 protein treated group at 2 dpi (Fig. 2F). As expected, footpad immune cells from CCL4-treated mice had a higher percentage of inflammatory monocytes, dendritic cells, macrophages, and NK cells than PBS treated controls, suggesting CCL4 treatment induced functional changes in immune cell mobilization (Supplementary Fig. 1). Overall, these data suggest that CCL4 depletion nor CCL4 protein inoculation has a significant impact on acute Mayaro disease or replication.

### CCR5 ligands are upregulated in response to MAYV infection, and lack of CCR5 does not impact Mayaro disease severity

CCL4 signals through the G protein-coupled receptor CCR5 (27). Lack of CCL4 or protein injection from external sources showed no significant impact on disease outcome (Fig. 2). Thus, we next tested whether genetic deletion of CCL4’s receptor, CCR5, would impact disease outcomes following MAYV infection. CCR5 also binds to other ligands, such as CCL3 and CCL5, which are also upregulated during MAYV infection (22, 24). First, to validate these previous reports, we evaluated the impact of MAYV infection on CCL3 and CCL5 at 2 dpi. Like CCL4, CCL3 and CCL5 were significantly upregulated following MAYV infection compared to mock-infected animals (Fig. 3A-B). Next, we infected WT and CCR5^-/-^ mice with MAYV and monitored for disease development and virus replication. We observed no differences in weight loss following infection (Fig. 3C), and similar footpad swelling and viremia at 2 dpi were observed in both groups (Fig. 3D-E). Overall, this data demonstrates that CCR5 does not significantly affect MAYV disease outcomes in mice.

**Figure 3.**
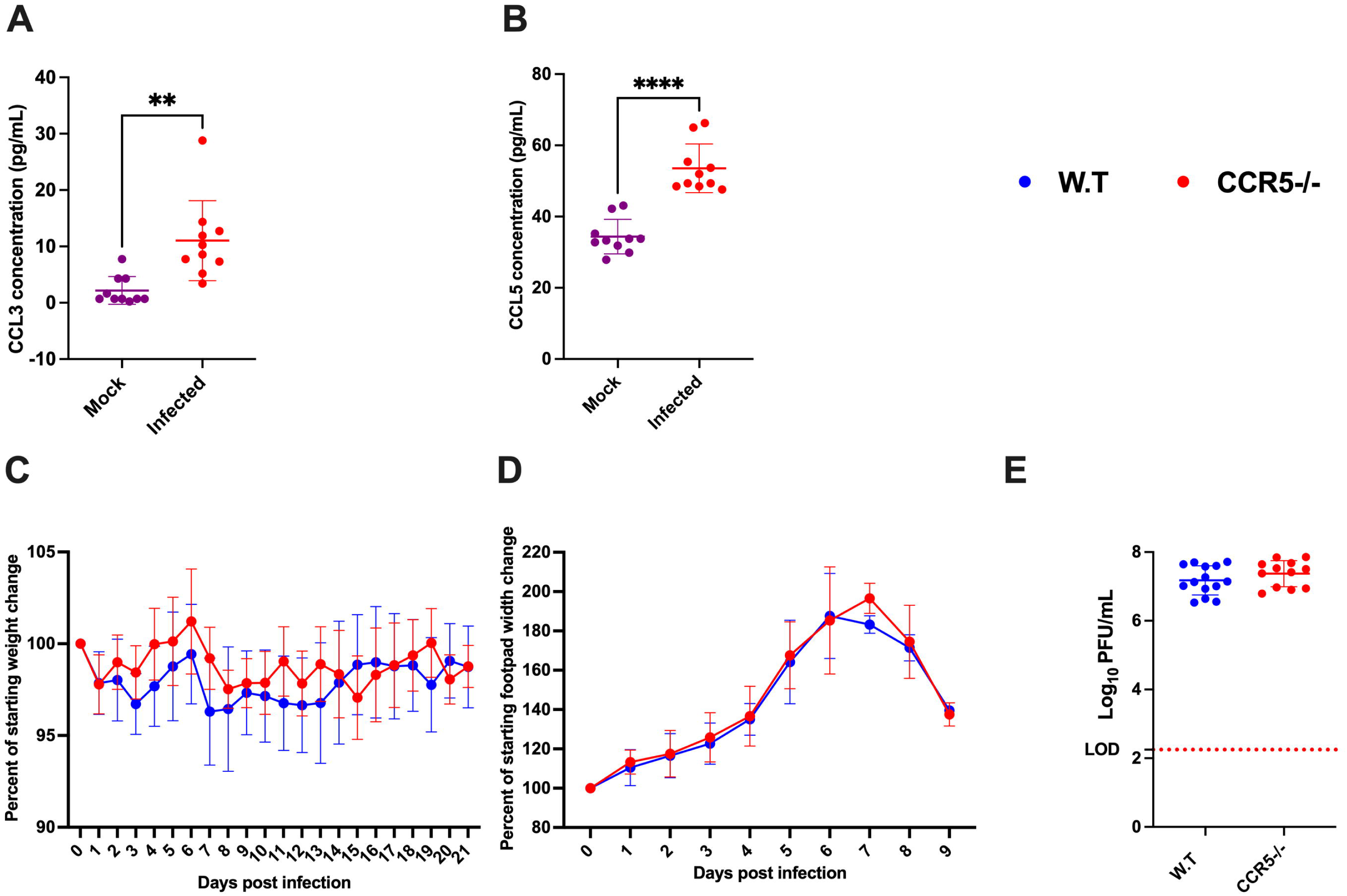
CCR5 ligands are upregulated in response to MAYV infection, and CCR5 depletion showed minimal impacts on Mayaro disease. A-B) WT mice were inoculated with 10^4^ PFU of MAYV strain TRVL 4675 through injection of both hind footpads. Serum samples were collected at 2 dpi to measure CCL3 (A) and CCL5 (B) by Luminex assay. Data are presented as picograms per milliliter of serum; unpaired t-test with Welch’s correction, **p < 0.01, ****p < 0.0001. C-E. WT and CCR5^-/-^ mice were inoculated with MAYV 10^4^ PFU/feet in both hind feet and monitored for disease development until 21 dpi. Weight loss (C), footpad swelling (D), and virus titer (E) was measured. Weight loss and footpad swelling were analyzed using multiple unpaired t-tests with the Holm-Sidak method for multiple comparisons, and viremia data was analyzed by an unpaired t-test with Welch’s correction. The error bars represent the standard deviation, bars indicate mean values, and the dotted line represents the limit of detection; (three experiments, n=13-14/group).

### CCR5 deletion showed minimal effect on peripheral or footpad immune cell profiles during MAYV infection

Next, we aimed to explore CCR5’s impact on the immune cell profile during MAYV infection. We isolated blood leukocytes at 2 and 6 dpi and performed flow cytometry. At 2 dpi, we observed similar percentages of CD4 T cells (Fig. 4A), neutrophils (Fig. 4C), inflammatory monocytes (Fig. 4D), and dendritic cells (Fig. 4E) in both WT and CCR5^-/-^ groups. Notably, CD8 T cells were reduced significantly in CCR5^-/-^ group compared to WT animals (Fig. 4B). At 6 dpi, no differences were observed in any of the tested immune cell types (Supplementary Fig. 2). We also assessed footpad immune cells at 6 dpi at peak footpad swelling. We found similar levels of CD4 (Fig. 5A) and CD8 T cells (Fig. 5B), neutrophils (Fig. 5C), inflammatory monocytes (Fig. 5D), dendritic cells (Fig. 5E), macrophages (Fig. 5F), and NK cells (Fig. 5G) between CCR5^-/-^ and WT group. Altogether, these data highlight that CCR5 has minimal impacts on immune cell profiles during MAYV infection.

**Figure 4.**
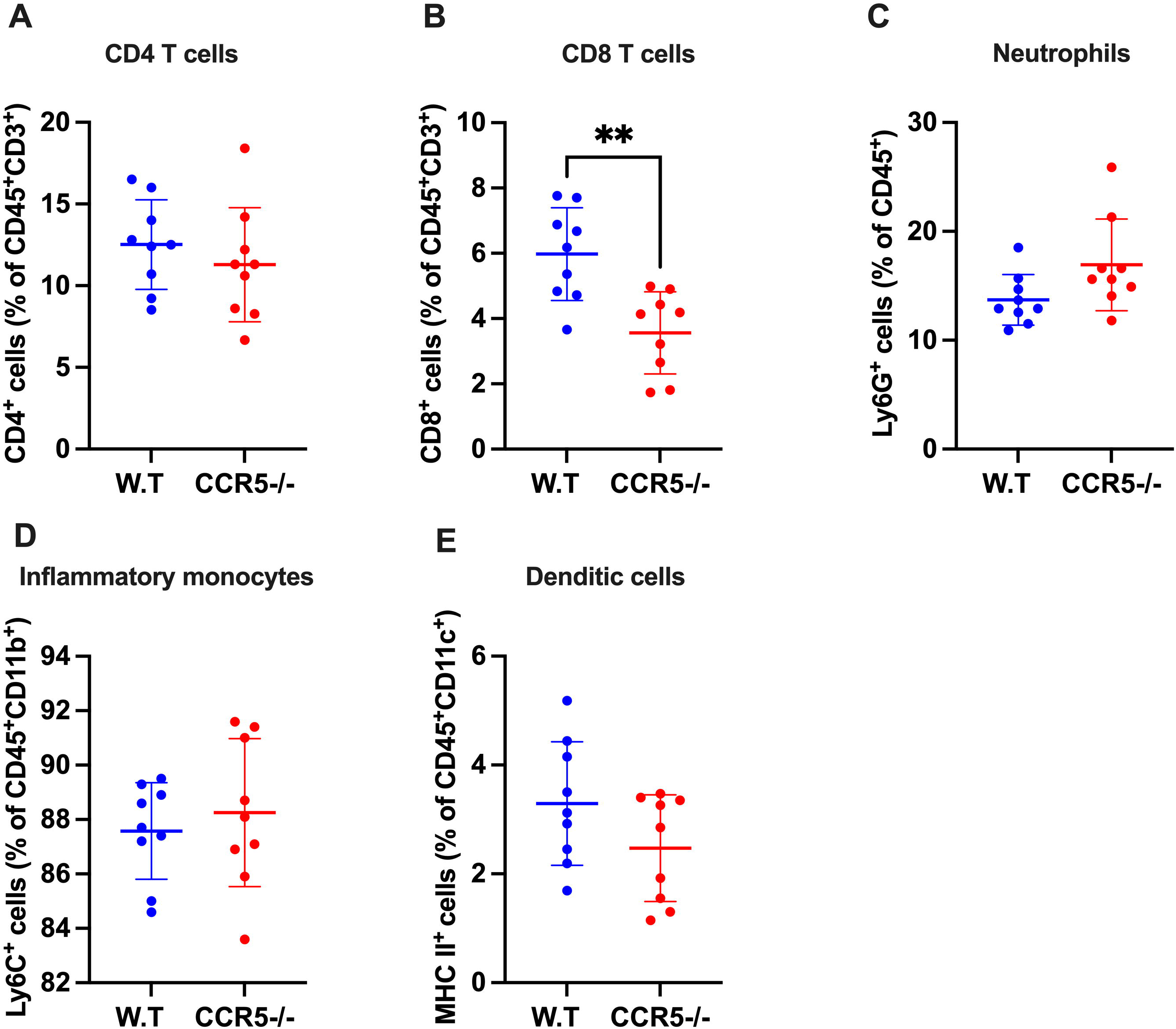
CCR5 deletion showed reduced CD8 T cell populations in blood during peak viremia. WT and CCR5^-/-^ mice were inoculated with 10^4^ PFU of MAYV in both hind feet, and blood was collected at peak viremia (2 dpi) to determine the immune cell population. A-E Plots presenting the percentage of CD4 T cells (CD45^+^CD3^+^CD4^+^) (A), CD8 T cells (CD45^+^CD3+CD4^+^) (B), neutrophils (CD45^+^Ly6G^+^) (C), inflammatory monocytes (CD45^+^CD11b^+^Ly6C^+^) (D), and dendritic cells (CD45^+^CD11c^+^MHCII^+^) (E). Immune cell percentage data was analyzed with an unpaired t-test with Welch’s correction. The error bars represent the standard deviation, bars indicate mean values, and asterisks indicate statistical differences; **p < 0.001, two experiments, n=9/group.

**Figure 5.**
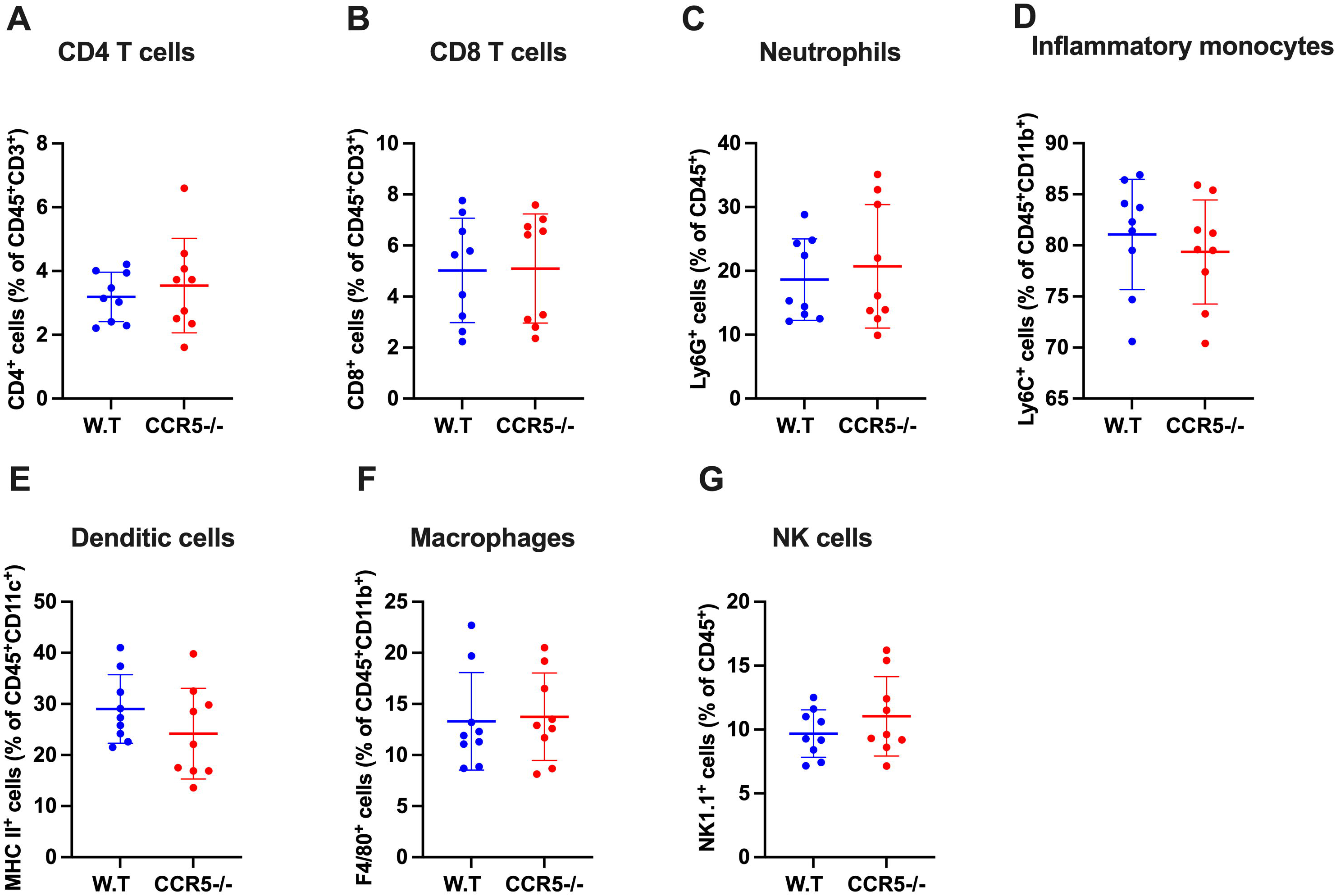
CCR5 deficiency showed no effect on immune cell populations in footpads during peak footpad swelling. WT and CCR5^-/-^ mice were inoculated with 10^4^ PFU of MAYV in both hind feet, and footpads were collected at 6 dpi to assess immune cell populations. A-G Plots presenting the percentage of CD4 T cells (CD45^+^CD3^+^CD4^+^) (A), CD8 T cells (CD45^+^CD3^+^CD4^+^) (B), neutrophils (CD45^+^Ly6G^+^) (C), inflammatory monocytes (CD45^+^CD11b^+^Ly6C^+^) (D), dendritic cells (CD45^+^CD11c^+^MHCII^+^) (E), macrophages (CD45^+^CD11b^+^F4/80^+^) (F), and NK cells (CD45^+^NK1.1^+^) (G). Immune cell percentage data was analyzed with an unpaired t-test with Welch’s correction. The error bars represent the standard deviation and bars indicate mean values; two experiments, n=9/group.

## Discussion

MAYV infection causes acute and chronic disease characterized by fever, skin rash, myalgia, and debilitating joint pain. In infected mammalian hosts, immune cells migrate to target tissues such as muscles, joints, and synovial tissues, leading to the initiation of inflammatory response and up-regulation of several proinflammatory cytokines and chemokines, including CCL4 (19, 42). Here, we explored the role of CCL4 in Mayaro disease. We depleted CCL4 through antibody-based neutralization and injected recombinant CCL4 protein and observed no effect on disease outcomes or viral replication. Furthermore, we infected mice deficient in CCL4’s receptor, CCR5, and found minimal impact on disease development, likely indicating that CCL4 and CCR5 play a minimal role in MAYV disease.

CCL4 is produced by various cell types, including monocytes, macrophages, natural killer cells, neutrophils, B and T lymphocytes, fibroblasts, and stromal cells (43–49). Plasma samples of humans infected with arthritogenic alphaviruses such as CHIKV, RRV, and MAYV have high levels of pro-inflammatory cytokines and chemokines, including CCL4 (15–19, 50). This highlights that CCL4 is broadly upregulated in response to different arthritogenic alphaviruses. Therefore, we hypothesized that CCL4 contributes to arthritogenic alphavirus disease. To test this, we depleted CCL4 using a neutralizing antibody, treated mice with recombinant CCL4, and used mice deficient in CCR5 followed by infection with MAYV. Following CCL4 depletion or treatment with CCL4 protein, we observed no difference in disease development between the groups. CCL4’s receptor, CCR5, is also used by other chemokines CCL3, CCL5, CCL8, CCL11, and CCL3L1 (24, 27, 51–53); as such, we hypothesized that compensation by the other ligands may have obscured impacts on disease outcomes and thus tested CCR5^-/-^ mice. CCR5 is expressed on T cells, dendritic cells, macrophages, and eosinophils, and ligands initiate chemotaxis to the site of infection (24, 27, 51, 54). We observed that CCR5 ligands CCL3, CCL4, and CCL5 were upregulated in MAYV-infected animals at 2 dpi. Surprisingly, when we infected CCR5^-/-^ animals, we observed only minor or no differences in disease phenotypes, viral replication, and immune cell populations compared to WT controls. One possible explanation is that lack of CCR5 might lead to a stronger interaction between CCL3, CCL4, and CCL5 to other receptors such as CCR1, CCR3, CXCR3, or CCR4 for compensatory immune cell activation (55, 56). Given the redundancy with other chemokine receptors, it is possible that other receptors compensate for the loss of CCR5 in mice infected with MAYV (57). The impact of other receptors such as CXCR3, CCR1, and CCR3, among others, on arthritogenic alphavirus disease outcomes should be explored in future investigations. Moreover, given the redundancy of the chemokine/receptor systems, mice with deletions in several ligands or multiple receptors may shed light on the role of these chemokines (58) in alphavirus disease. Furthermore, these chemokines can be neutralized together using monoclonal antibodies.

There are several key immune cells that express CCR5 and contribute either positively or negatively to alphavirus disease and replication, including monocytes, macrophages, NK cells, γδ T cells, CD8^+^ and CD4^+^ T cells, and B cells (40). After alphavirus infection, chemokines like CCL4 are secreted, recruiting immune cells to the site of infection (59, 60). The monocytes promote local virus replication and enhance the transport of the virus to distal sites (61, 62). Additionally, alphaviruses can replicate in macrophages (62, 63), which act as reservoirs for viral RNA in infected tissues (64). Beyond viral replication and dissemination, tissue-resident myeloid cells also produce interferons and pro-inflammatory cytokines that aid in viral clearance yet also promote inflammatory tissue damage (65, 66). A pathogenic role of NK cells has been proposed during arthritogenic alphavirus infection (67). CHIKV infection in mouse footpads leads to an increase in γδ T cells in the foot (68), where they play a protective role. CD8^+^ T cells may also exert an antiviral effect during infections caused by arthritogenic and neurotropic alphaviruses (69–72). However, arthritogenic alphaviruses can evade the CD8^+^ T cell response (69, 73) which may partly explain why infection persists in joint-associated tissues (74, 75). Mice with genetic or acquired deficiencies of CD4^+^ T cells developed minimal or no joint pathology when challenged with CHIKV, suggesting a pathogenic role in disease (76). During alphavirus infection, B cells produce virus-specific antibodies that clear virus from the bloodstream (77). Infection of B cell-deficient mice with CHIKV resulted in persistent infection in the joint (77). Considering the important role of chemokines and their receptors in the recruitment of immune cells for alphavirus infection control, future studies should be conducted by neutralizing the redundant chemokines together with antibodies or knockout of redundant receptors in mouse models.

Despite exerting minimal impacts on Mayaro infection and disease progression, CCR5—and most likely its ligands—contribute to the development of other viral diseases. For example, mice deficient in CCR5 are fully protected from disease following infection with a mouse-adapted dengue virus (78). Similarly, mice treated with Met-RANTES (Met-R), a CCR5 inhibitor, were similarly protected against disease and DENV replication. In contrast, CCR5^-/-^ mice infected with Japanese encephalitis virus (JEV) or West Nile virus (WNV) have significantly worse disease outcomes and skewed immune cell profiles (79, 80). Similarly, CCR5-deficient mice infected with influenza virus have increased mortality rates (81), suggesting a complex role for CCR5 in viral infection. Like in influenza virus infection, CCL5^-/-^ and CCR5^-/-^ had worse disease outcomes than WT mice, which was associated with virus-induced apoptosis (82). Surprisingly, CCL3 deficient mice were protected from lung inflammation compared to WT controls when infected with influenza virus (83), suggesting a complex interplay between various chemokines and their receptors during influenza virus infection. Finally, antibody depletion of CCL3 resulted in worse disease outcomes and altered immune cell activation following respiratory syncytial virus infection (84). Taken together, these data further underscore the complexity and redundancy of mammalian chemokine signaling and highlight the importance of virus-specific impacts for each ligand and/or receptor.

In summary, our study provides insights into the role of CCL4 and CCR5 in MAYV pathogenesis. The results suggest that CCL4 nor CCR5 significantly alter disease outcomes, viral replication, or immune cell populations. Therefore, CCL4 or CCR5-based therapeutics may not be effective for arthritogenic alphavirus disease. However, given the redundancy of mammalian chemokine signaling, future studies should explore the role of other chemokine receptors individually or in conjunction with CCR5.

## Acknowledgements

We are grateful to Melissa Makris for assisting with flow cytometry analysis. This work was supported by NIAID R21AI153919-01 awarded to J.W-L.

